# Parameterizing Pantherinae: *de novo* mutation rate estimates from *Panthera* and *Neofelis* pedigrees

**DOI:** 10.1101/2024.04.06.587788

**Authors:** Ellie E. Armstrong, Sarah B. Carey, Alex Harkess, Gabriele Zenato Lazzari, Katherine A. Solari, Jesús E. Maldonado, Robert C. Fleischer, Neel Aziz, Patricia Walsh, Klaus-Peter Koepfli, Eduardo Eizirik, Dmitri A. Petrov, Michael G. Campana

**Author notes:** designates equal contribution (Dmitri A. Petrov, Michael G. Campana).

## Abstract

Estimates of *de novo* mutation rates are essential for phylogenetic and demographic analyses, but their inference has previously been impeded by high error rates in sequence data and uncertainty in the fossil record. Here, we directly estimate *de novo* germline mutation rates for all extant members of *Panthera,* as well as the closely related outgroup *Neofelis nebulosa*, using pedigrees. We use a previously validated pipeline (RatesTools) to calculate mutation rate for each species and subsequently explore the impacts of the novel rates on historic effective population size estimates in each of these charismatic felids of conservation concern. Importantly, we find that the choice of reference genome, the data type and coverage, and the individual all impact estimates of the mutation rate. Despite these stochastic effects, we inferred that base pair mutation rates for all species fell between 0.5 and 1.4e-08 per generation per base pair (mean 0.81e-08 ± 0.35-08 across Pantherinae). Our results provide a cautionary view on inter-species mutation rate comparisons, given the error associated with the reference genome choice and sequencing depth of coverage of the individuals.

## Introduction

The germline *de novo* mutation rate can be estimated in several ways, including indirectly through phylogenetic methods, directly via observation of *de novo* mutations in trios, pedigrees, or germline tissue, or through mutation accumulation experiments. For organisms that cannot be manipulated in the laboratory, indirect phylogenetic methods and direct estimates using pedigrees or germline tissue are the only feasible options. Until very recently, the inference of neutral mutation rates using indirect methods dominated the field. This choice of method was primarily driven by sequencing error rates exceeding most species’ mutation rates, thereby making the discovery and validation of real *de novo* mutations via direct methods exceedingly difficult. However, improvements in sequencing technologies have recently rendered direct estimation more feasible (Pfeifer 2017; Koch et al. 2019; Bergeron et al. 2022; Bergeron et al. 2023).

The neutral substitution rate can be estimated by counting the number of fixed, putatively neutral mutations between lineages, overlaid with assumed divergence times that are often calibrated with fossil evidence. Estimates of neutral substitution rates have traditionally relied on the assumption that mutations accumulate at a steady rate, also known as the “molecular clock hypothesis” (Zuckerkandl 1962). This assumption leads to the idea that the rate of neutral sequence divergence is equivalent to the substitution rate per year (Kimura 1968), although these assumptions can be relaxed (Lepage et al. 2007). While the substitution rate combines effects of both mutation and fixation, these rates are frequently used as an estimate of the mutation rate parameter (e.g. Ho and Larson 2006). Neutral substitution/mutation estimates from divergence can be error-prone due to differences in the way orthologous regions are aligned, decisions regarding which regions to include in the analysis, variable interpretations or the availability of fossil evidence, uncertainty in the phylogenetic placement of fossil calibration priors, and the unaccounted shifts in the rate of the molecular clock along the path of divergence. In estimating the divergence between closely related species, a challenge arises in determining which observed differences between lineages are fixed rather than still segregating, especially since neutral rates are often estimated using one representative individual per lineage. Indirect estimates of mutation rate using these phylogenetic methods have shown inconsistencies that might be partly attributed to the employment of different algorithms and methods (Shendure and Akey 2015; Bergeron et al. 2022). Furthermore, indirect inferences have been shown to differ substantially from the rates estimated directly from pedigrees (Kong et al. 2012; Scally and Durbin 2012; Moorjani et al. 2016; Zhang et al. 2022).

In the conservation genetics literature, projection of past population sizes has played an important role in our understanding of species’ current distributions and diversity, and how these are shaped by climatic and geologic events, as well as the influence of recent anthropogenic perturbations (e.g., Wilder et al. 2023). The mutation rate is commonly used to scale results from methods that use the coalescent to infer historic effective population size (N_e_) from genomic data (Li and Durbin 2011; Terhorst et al. 2017; Schiffels and Wang 2020). It has been well-documented that differences in the scaling parameters (mutation rate, recombination rate, and generation time) of these methods can cause vastly different projections of past population sizes (Nadachowska-Brzyska et al. 2016; Beichman et al. 2018; Armstrong et al. 2020; Zhang et al. 2022) and timing of demographic events (Campana et al. 2020). Despite this, neutral mutation rates inferred using phylogenetic approaches or a blanket ‘mammalian’ rate of 1.0e-09 per base pair per generation or 2.22e-09 per base pair per year (Kumar and Subramanian 2002) are most commonly applied in non-model mammalian species (Mather et al. 2020). Estimates of past effective population sizes are often interpreted without caveat and overlaid with other historic events to create a narrative that fits the data, but also may be wrought with error.

Mutation rates can be estimated from trios under the assumption that novel mutations can be identified by comparing the parental genotypes to offspring genotypes. Analyses are most commonly restricted to sites which are homozygous in the parents and heterozygous in the offspring. Direct estimates from pedigrees have become more common over the last several years as sequence quality has improved and estimates have been made for chimpanzees (Besenbacher et al. 1999), green monkeys (Pfeifer 2017), bears (Wang et al. 2022), wolves (Koch et al. 2019), birds (Smeds et al. 2016; H. Zhang et al. 2023), whales (Suárez-Menéndez et al. 2023), fish (C. Zhang et al. 2023), and bees (Liu et al. 2017). Most recently, Bergeron et al. (2023) estimated germline mutation rates across 68 species of fishes, reptiles, birds, and mammals using high-quality genome data from 151 trios, including a trio each for tiger and leopard. Importantly, the inference of *de novo* mutation rates from pedigree data has suggested that mutation rates are variable across species, and that, even within mammals, mutation rates differ by a factor of 40 (Bergeron et al. 2023)

The felid genera *Panthera* and *Neofelis,* which together comprise the subfamily Pantherinae, represent one of the most popular and charismatic groups of species in the world. Most members of these genera are categorized between ‘Near Threatened’ and ‘Endangered’ by the IUCN Red List of Threatened Species (https://www.iucnredlist.org). There has been significant discussion in the scientific literature regarding the timing and order of the divergence of species in the *Panthera*-*Neofelis* clade. Recent estimates suggest that the split between *Panthera* and *Neofelis* occurred roughly between 4 and 6 million years ago (Li et al. 2016; Bursell et al. 2022; Yuan et al. 2023). However, pinpointing the exact timing of these divergence events and resolving the relationships among the species have proven difficult. This is in large part due to potential historic hybridization and/or introgression among species or extensive lineage sorting due to a very fast sequence of speciation events (Li et al. 2016; Figueiró et al. 2017; Li et al. 2019). The clade has been particularly difficult to date due to the lack of fossil evidence and the inability to distinguish fossil *Panthera* specimens due to their phenotypic similarities (Christiansen 2008; Tseng et al. 2014). In addition, as apex predators, their naturally small population sizes have put many populations and entire species at risk of extinction. Accurately timing demographic events such as bottlenecks and expansions, as well as resolving the timing of possible introgression and hybridization events amongst species, hinges on an accurate inference of the mutation rate.

There have been several studies that have provided estimates of neutral mutation rates using indirect methods in *Panthera* and *Neofelis (Cho et al. 2013; Kim et al. 2016; Liu et al. 2018)*, as well as recent direct estimates for the tiger and leopard (Bergeron et al. 2023). These rate estimates span from 1.1e-09 to 1.0e-08 per base pair per generation, covering nearly an order of magnitude difference (Figure 1; Table S1). Such a large uncertainty in these estimates significantly impacts the inferences drawn about historic population sizes.

**Figure 1:**
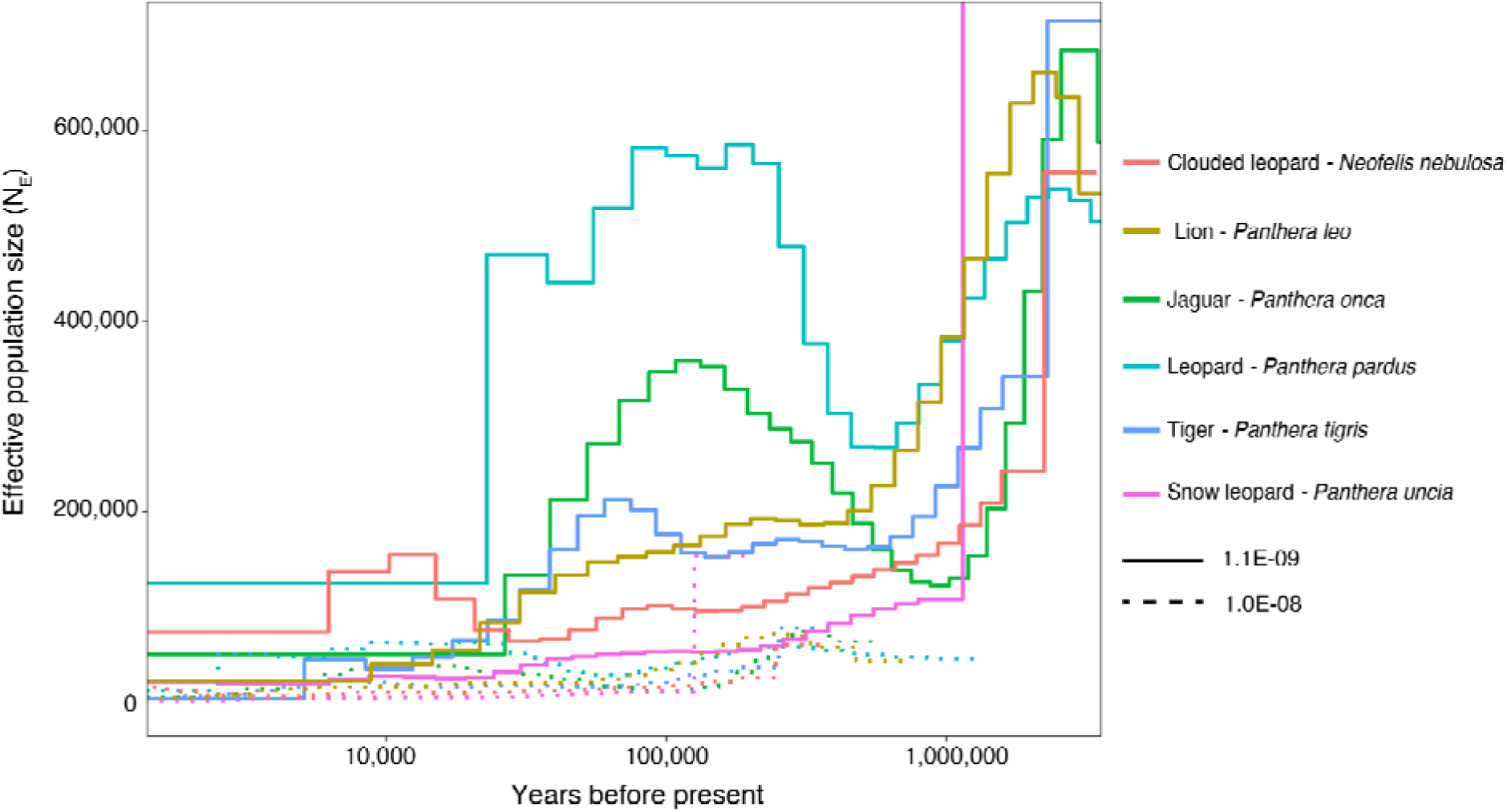
PSMC plot for *Panthera* spp. and *Neofelis nebulosa* showing N_e_ inferred when using previously estimated maximum (1.0e-08) and minimum (1.1e-09) per-generation mutation rates.

Here, we use the RatesTools pipeline (Armstrong and Campana 2023) to establish germline mutation rates for species of *Panthera* and *Neofelis* using direct estimates from novel whole-genome pedigree data from the tiger (*Panthera tigris*), lion (*Panthera leo*), snow leopard (*Panthera uncia*), jaguar (*Panthera onca*), and the mainland clouded leopard (*Neofelis nebulosa*), as well as from existing data with previously estimated rates from a tiger and leopard (*Panthera pardus*) trio (Bergeron et al. 2023). We also investigate the impact of these novel rates on predicting historic effective population size (N_e_) and assess the impact of reference genome choice and data type on mutation rate inference. We show that extant Pantherinae have broadly similar mutation rates across the clade (between 0.5 and 1.4e-08 per base pair per generation, mean 0.81e-08 ± 0.35-08), with trio-inferred rates generally falling near the higher (1.0e-8) bound of previously employed rates. Moreover, we demonstrate that reference genome choice, the employed data type(s) and sequencing depth, and the assayed individuals included in the trios, all impact estimates of the mutation rate.

## Materials & Methods

### Sample collection and sequencing

Details of newly sequenced *Panthera* and *Neofelis* trios are provided in Table S2 and Figure S1. Sequencing procedures are detailed below. We additionally downloaded sequence read data from NCBI Sequence Read Archive for the tiger and leopard trios generated as part of (Bergeron et al. 2023) for analysis (tiger: accessions SRR17072401, SRR17072410–17, SRR17072419–22, SRR17072424–25; leopard: accessions SRR17072427, SRR17072429, SRR17072432–37, SRR17072707).

### Lions

Lion samples were provided by the Smithsonian’s National Zoo and Conservation Biology Institute (NZCBI) from lions “Luke” (sire, SB114161), “Nababiep” (dam, SB114162), and two of their male offspring from the same litter (“Aslan”, SB114626 and “Baruti”, SB114625) (Table S2). Aslan and Baruti were transferred to the Calgary Zoo in 2012. EDTA whole-blood samples were opportunistically collected from each individual during routine veterinary check-ups and subsequently frozen at -80°C prior to shipping and processing. Additionally, previously extracted DNA samples for Luke and Nababiep (extractions used the Qiagen Blood and Tissue Kit [Qiagen, Inc., Germantown, MD] using a Qiagen BioSprint® 96 robot according to manufacturer’s instructions) were built into dual-indexed Illumina libraries using a modified Illumina Nextera (Illumina, Inc., San Diego, CA) DNA library preparation protocol as described in (Baym et al. 2015) and 2 × 150 base pair (bp) paired-end sequenced at Admera Health (South Plainfield, NJ) on a NovaSeq 6000 S4 lane (Illumina, Inc., San Diego, CA).

Blood samples from Aslan and Baruti (100 µl whole blood) were extracted using the Qiagen DNeasy Blood and Tissue Kit. Provided instructions were followed, with the exception that AE buffer was used instead of phosphate buffered saline during the digestion step and the total volume during digestion was 220 µl. Extracted DNA samples were sent to Admera Health (South Plainfield, NJ) and were prepared using the KAPA HyperPrep kit (Kapa Biosystems, Inc., Wilmington, MD) according to the provided instructions. Samples were 2 × 150 bp paired-end sequenced on one lane each of a HiSeq X Ten instrument (Illumina, Inc., San Diego, CA). Samples for Luke and Nababiep were prepared by HudsonAlpha Institute for Biotechnology (Huntsville, AL) for 10x Genomics Chromium sequencing using 100 µl whole blood samples from each individual. Samples were extracted using the MagAttract HMW DNA Kit (Cat#:67563; Qiagen, Inc., Germantown, MD) and Chromium libraries were prepared according to provided instructions for all of the samples. The libraries were then shipped to Admera Health and were paired-end sequenced (2 × 150 bp) on a lane of a HiSeq X Ten instrument.

### Tigers

Tiger samples were provided by In-Sync Exotics (Wylie, TX) from three tigers “Assad” (sire), “Zahra” (dam), and their male offspring “Kylo-Ren”. EDTA whole-blood samples were taken during routine veterinary check-ups. Samples for “Zahra” and “Assad” were aliquoted into 5 mL volumes for 10x Genomics Chromium sequencing and shipped on ice to HudsonAlpha (Huntsville, AL). Samples were prepared and sequenced as described above. Kylo-Ren’s 10x Genomics Chromium sequencing data were previously reported in (Armstrong et al. 2022; Armstrong et al. 2024) (NCBI accession SRR16296779, ). Additionally, standard paired-end libraries for all individuals were prepared using the Illumina Nextera DNA library preparation protocol with adjustments as described in (Baym et al. 2015). Nextera libraries were shipped to Admera Health (South Plainfield, NJ) and sequenced on a Hiseq X Ten instrument.

### Snow leopards

EDTA whole-blood samples from four snow leopards “Raj” (sire; SB117026), “Anna” (dam; SB117027), their female offspring “Tikka” (SB118001), and male offspring “Tsering” (SB118009) were provided by San Francisco Zoo and were collected as part of routine veterinary care. Samples were extracted using a Qiagen DNeasy Blood and Tissue Kit according to the provided instructions. Genomic libraries were prepared using the same modified Illumina Nextera DNA library preparation protocol cited above (Baym et al. 2015). The libraries were then shipped to Admera Health and were paired-end sequenced (2 × 150 bp) on a lane of the NovaSeq X instrument (Illumina, Inc., San Diego, CA).

### Jaguars

We analyzed whole-genome sequences from nine jaguar individuals that comprised an extended pedigree containing four trios (Figure S1). These animals were sampled during routine veterinary check-ups as part of an Argentinian *ex-situ* breeding and rewilding program. Their genomes were paired-end sequenced (2 × 150 bp) on an Illumina NovaSeq X platform (Illumina, Inc., San Diego, CA) as part of a project that developed a SNP panel for forensic and molecular ecology analyses of this species (Lazzari et al., *in prep*).

### Mainland clouded leopards

Approximately 3 ml of whole blood were collected into EDTA Vacutainer tubes (Becton Dickinson, Franklin Lakes, NJ) during routine veterinary exams from three mainland clouded leopards maintained in the living collection of NZCBI: “Hannibal” SB1278 (sire), “Jao Chu” SB1276 (dam), and “Ta Moon” SB1433 (male offspring) (Table S2). The samples were delivered to Psomagen, Inc. (Rockville, MD) for DNA extraction, library preparation, and whole genome sequencing. Genomic DNA was extracted from a 300 µl aliquot of whole blood using the Mag-Bind Blood and Tissue Kit (Omega Bio-Tek Inc., Norcross, GA). DNA concentration was evaluated with Picogreen and Victor X2 fluorometry (Life Technologies, Carlsbad, CA), and an Agilent 4200 Tapestation (Agilent Technologies, Santa Clara, CA), and quality was checked via electrophoresis on a 1% agarose gel. Extracted DNA was then sheared and enriched into 350 bp fragments using ultrasonication (Covaris S220, Woburn, MA), which were used to construct a genomic library using the TruSeq DNA PCR-free library kit (Illumina, San Diego, CA). Library quality was assessed with an Agilent 4200 Tapestation and Lightcycler quantitative PCR (Roche Life Science, St. Louis, MO) and then paired-end sequenced (2 × 150 bp) to a minimum depth of 20x on an Illumina NovaSeq 6000 instrument. The genome of an additional offspring (“Sa Ming”, SB1434, brother of “Ta Moon”) was previously reported in Bursell et al. 2022 (NCBI accession SRR13774417) and included in the analyses.

### Lion Genome Assembly

Pathology tissues from the post-mortem autopsy, including liver (160 mg), heart (280 mg), spleen (190 mg), and whole blood (350 µl) from lion “Luke” were shipped on dry ice to HudsonAlpha (Huntsville, AL). DNA was extracted using the PacBio Nanobind Tissue Kit (SKU 102-302-100) according to standard instructions, with 350 µl of whole blood as input. The SMRTbell Prep Kit 3.0 was then used to prepare a SMRT library for Pacific Biosciences HiFi sequencing and sequenced across two SMRT cells on a PacBio Revio system. 60 mg of heart tissue was used as input for the Dovetail Omni-C Kit (CAT#21005) and a Hi-C library prepared according to standard procedures for tissue processing provided in the Dovetail Omni-C User Guide 2.0. The Omni-C library was sequenced to approximately 30× coverage on an Illumina NovaSeq 6000 instrument (Illumina, Inc., San Diego, CA).

We first used resequence data (Armstrong et al. Unpublished data) from 12 lions (6 females, 6 males; Table S6) in order to detect kmers unique to males (henceforth known as y-mers). We filtered the data using Trimmomatic v0.39 with leading and trailing values of 3, sliding window of 30, jump of 10, and a minimum remaining read length of 40 (Bolger et al. 2014) and identified k-mers using meryl v1.3 *count* (Rhie et al. 2020). We used meryl’s *intersect* function to identify all 21-mers shared in females, followed by *difference* to identify the 21-mers only found in males.

We created an initial assembly using HiFiasm v0.19.6 (Cheng et al. 2021) as a Hi-C integrated assembly by providing the Pacbio HiFi reads, as well as the Dovetail Omni-C reads using the ‘--h1’ and ‘--h2’ flags. To identify and correctly phase the XY sex chromosomes, we used a two-step approach (Carey et al., *in prep*). First, Y-mers were mapped to the assemblies in order to identify putatively Y-associated contigs using BWA-MEM v0.7.17 (Li and Durbin 2009; Li 2013) with flags ‘-k 21’, ‘-T 21’, ‘-a’, and ‘-c 10’. Next, we used the YaHS program v1.1a-r3 (Zhou et al. 2022) in order to scaffold a concatenated assembly. The goal of this step was to manually inspect the assembly in Juicebox in order to identify the putative X and Y chromosome scaffolds (ignoring the autosomes). This is most noticeable because of the ‘X’ (overlapping diagonal and off-diagonal) pattern created in the Hi-C heatmap by the pseudoautosomal region (PAR), which is the only region that recombines on the X and Y chromosomes. The distinct haplotypes produced by HiFiasm were concatenated and Hi-C data mapped according to Phase Genomics suggestions (https://phasegenomics.github.io/2019/09/19/hic-alignment-and-qc.html). Briefly, the concatenated assembly was indexed using BWA v0.7.17 *index* with flags ‘-a bwtsw’ (Li and Durbin 2009). Omni-C reads were mapped using BWA-MEM using the ‘-5SP’ flag (Li 2013). Reads were then marked for PCR duplicates using SAMBLASTER v0.1.26 (Faust and Hall 2014) and converted to BAM format using SAMtools v1.16.1 *view* with flags ‘-S’, ‘-h’, ‘-b’, and ‘-F 2304’ (Li et al. 2009; Danecek et al. 2021). The haplotype combined fasta was indexed using SAMtools *faidx.* Next, we used Juicer Tools v1.9.9 in order to convert our file for modification and visualization in Juicebox (Durand et al. 2016). A contig was considered Y-linked if Y-mer coverage was >1 and/or it scaffolded to other Y-linked contigs. A total of 67 scaffolds were putatively identified as being Y-linked, while 9 were identified as being X-linked (See *Supplementary Information* for details). Putative Y scaffolds were all moved into haplotype one from the original, non-concatenated HiFi assemblies, while putative X scaffolds were all moved into haplotype two. The result of this step is theoretically two distinct haplotypes with a complete set of autosomes and either all the putative X or Y scaffolds.

The distinct haplotypes were then re-scaffolded separately using the same approach as above (Omni-C read mapping, YaHS scaffolding). Both haplotypes were visualized in Juicebox and manual edits were made. Specifically, we broke apart one scaffold in each haplotype where there were telomere misjoins and moved several other fragments based on the contact map visualization (See *Supplementary Information* for details). We then checked for contamination in each assembly using FCS-GX v1.2.4-1.el7 (Astashyn et al. 2023) and subsequently investigated the orientation and presence of telomere motifs using the GENESPACE v1.3.1 (Lovell et al. 2022) program. Debris occurring at the end of scaffolds after telomere motifs was removed and scaffolds were oriented and ordered according to the domestic cat genome (GCA_000181335.6; Buckley et al. 2020).

After manual curation, we calculated assembly statistics using Assemblathon2 scripts (Bradnam et al. 2013). We evaluated gene completeness using compleasm v0.2.2 (Huang and Li 2023) with the carnivora_odb10 library from BUSCO (Simão et al. 2015).

### De Novo Mutation Rate Analysis

We identified candidate *de novo* mutations (DNMs) using RatesTools 1.2.0 (Armstrong and Campana 2023) (see *Supplementary Information* for details regarding RatesTools upgrades since its initial publication). For 10x Genomics Chromium data, we trimmed the first 16 bp of read 1 (corresponding to the 10x Genomics Chromium barcode) using Seqtk 1.4 (Li 2023: https://github.com/lh3/seqtk) as we found that the barcode region negatively impacted the retained callable genome using the Genome Analysis Toolkit 4.4.0.0 (McKenna et al. 2010; Table S15). We then combined the trimmed 10x Genomics Chromium data with the standard Illumina libraries for downstream analysis (See *Supplementary Information* for analyses partitioned by library type). Afterwards, using RatesTools, we aligned reads to and called genotypes using the species-appropriate nuclear genome reference(s). Lion trios were mapped to the original PanLeo1.0 assembly (Armstrong et al. 2020) and the ‘mappable’ novel lion genome reported here (defined as the assembly’s haplotype 2 including X chromosome with the Y chromosome added). Tiger trios were mapped to the GenTig1.0 (Armstrong et al. 2022) and PanTigT.MC.v3 (GCA_021130815.1; Shukla et al. 2022) genomes. Clouded leopards were mapped to three genome assemblies: a short-read assembly (SaMing-1434 [GCA_027422525.1; Bursell et al. 2022]) and two long-read assemblies (SNNU_Nneo_1 [GCA_030324275.1; Yuan et al. 2023] and the primary haplotype assembly of mNeoNeb1 [GCA_028018385.1; Vertebrate Genomes Project 2023]). Leopard, snow leopard, and jaguar data were mapped to the most contiguous reference genome available for each respective species (leopard: Ppardus1 [GCA_024362965.1; Armstrong et al. 2022]; snow leopard: PanUnc1.0 [GCF_023721935.1; Armstrong et al. 2022]; jaguar: Panthera_onca_HiC [GCF_028533385.1; DNAzoo.org, Dudchenko et al. 2017; Dudchenko et al. 2018]. The draft assembly for the jaguar was generated by the DNA Zoo team from short insert-size PCR-free DNA-Seq data using w2rap-contigger (Clavijo et al. 2017); see (Dudchenko et al. 2018) for details. As described in Armstrong and Campana 2023, we annotated each genome’s 30,2- mappability using GenMap 1.2.0 (Pockrandt et al. 2020) and identified repetitive regions using RepeatMasker 4.1.5 (Smit, A., R. Hubley, and P. Green. 2013-2015)‘-gccalc -nolow -xsmall’; (Smit, A., R. Hubley, and P. Green. 2013-2015) with the *Felidae* repeat library and RepeatModeler 2.0.5 (Flynn et al. 2020) under default parameters.

RatesTools alignment, genotyping and variant filtration parameters were the same for all analyses except the per site maximum and minimum retained sequencing depths per individual, and minimum genotype quality (GQ) varied depending on the individual trio sequencing depths (Table S3). The per site max sequencing depth per individual was 125× for most analyses, except the maximum per site depth per individual was 250× for analyses in which individuals had sequencing depths greater than 50× (lion, tiger, and leopard trios). The minimum retained per site sequencing depth per individual was 20× for all analyses except for jaguar and clouded leopard, where we reduced the minimum retained depth to 10× due to the lower coverage (∼20× per individual) of these trios (See *Supplementary Information*). We retained sites with a minimum GQ of 65 per individual (Bergeron et al. 2023), except for trios in which the mean depth fell below 20× (the jaguar trios and one of the two *Neofelis* trios), where we reduced the minimum GQ to 45. We retained only sites confidently mapped to autosomes, except for the *Neofelis nebulosa* alignment against the scaffold-level SaMing-1434 assembly as these scaffolds are not anchored to chromosomes. Autosomal variants were phased using pedigree- phasing in WhatsHap 2.1 (Garg et al. 2016; Martin et al. 2016). We then filtered sites using VCFtools 0.1.16 (Danecek et al. 2011) (parameters: “--minDP <10 or 20> --minGQ <45 or 65> --maxDP <125 or 250> --max-missing 1 --min-alleles 1 --max-alleles 2“)and GATK 4.4.0.0 (parameters: ’--filter-name “filter” --filter-expression “QD < 4.0 || FS > 60.0 || MQ < 40.0 || SOR > 3.0 || ReadPosRankSum < 15 || MQRankSum < -2”’). We then removed low-mappability regions (GenMap mappability < 1.0), repetitive regions, and sites within 5 bp of an indel. We discarded scaffolds that were shorter than 100,000 bp before site- and region-filtering and those that were less than 10,000 bp after filtering. DNM candidates were identified with the calc_denovo_mutation_rate.rb parameters: “-b 100 -M 10 -w 100000 -l 100000 -S 50000 -- parhom --kochDNp --minAD1 --minAF 0.3”.

We then calculated initial mutation rates by dividing the number of candidate mutations by the total number of callable sites multiplied by two. After initial DNM candidate identification, we removed likely misalignments/complex mutations by removing any DNM candidate which had another candidate within 100 bp (based on typical alignment lengths of a single 150 bp read). We also removed any candidate DNM that appeared in multiple siblings. Afterwards, we calculated the final mutation rates by adjusting the initial rate estimates from calc_denovo_mutation_rate.rb by the fraction of sites that remained after removing likely misalignments and DNM candidates observed in siblings (See *Supplementary Information*). We then calculated 95% confidence intervals using the *binconf* function from the Hmisc 5.1-1 (Harrell et al. 2023) package in R 4.3.1 (R Core Team 2023). Complete parameter files for all runs are available in the Figshare repository (doi: 10.25573/data.25374244). We report mutation rates in mutations per base pair per generation.

We calculated average species rates by summing the total number of retained DNM candidates across all trios and dividing by the total summed callable genome across all trios multiplied by two. We then calculated binomial confidence intervals as above. Further, we calculated general Pantherinae-lineage-specific rates by calculating the mean, median, and standard deviation of the average species rates. For species where multiple reference genomes were analyzed, we only included the most contiguous (or curated) assembly (lion: novel genome assembly reported here; tiger: PanTigT.MC.v3; clouded leopard: mNeoNeb1).

### Demographic history

We next ran the Pairwise Sequential Markovian Coalescent (PSMC; Li and Durbin 2009) to estimate trajectories of historic N_e_. Though there are other options for reconstruction of demographic history, we selected PSMC because it is one of the only options for inferring past population sizes using single genomes, which is common in the field of conservation and evolutionary genetics. We ran 50 bootstrap replicates on high coverage, whole-genome data previously collected for lions, tigers, leopards, jaguars, snow leopards, and clouded leopards (Table S16). For each individual, in addition to the newly inferred rates, we also ran the minimum and maximum mutation rates established in previous research (Table S1).

Input PSMC files were generated by first indexing each species genome using BWA v0.7.17 (Li and Durbin 2009) *index*, flags ‘-a bwtsw’. Subsequently, each set of read pairs were then mapped to that species’ best representative genome (Table S16) using BWA-MEM (Li 2013) and sorted and indexed using SAMtools v1.16.1 (Danecek et al. 2021) *sort* and *index*, respectively. Duplicates were marked using Picard tools v2.18.14 (Broad Institute 2019). We followed the commands outlined here; https://github.com/lh3/psmc, with the exception that we replaced SAMtools with BCFtools v1.16 (Danecek et al. 2021) *mpileup* and removed the ‘-u’ flag. BCFtools *view* was additionally replaced with BCFtools *call*. No other changes were made to the pipeline in order to generate the PSMC and bootstrap files.

Results were visualized using the R packages psmcr v0.1-4 (https://github.com/emmanuelparadis/psmcr/) and tidyverse v1.3.2 (Wickham et al. 2019). We set the per generation mutation rate to the minimum and maximum as defined in Table S1. Generation time was set to five years for each lineage. Though we recognize these may vary based on the interpretation of generation time and various field data, we selected a generation time of five years to be consistent with previous publications which inferred demographic history using this method in the Pantherinae (Liu et al. 2018; Armstrong et al. 2020; Armstrong et al. 2021; Bursell et al. 2022; Solari et al. 2023). Bin size was set to 100. We additionally plotted the novel mutation rates defined by this study (Figure 3).

## Results

### Genome trio sequencing

Effective sequencing depths and breadths of coverage for each of the trio individuals analyzed here are provided in Tables S2 and S12. We sequenced the lion trio individuals to a mean of 46× (range: 37×–55×). The newly sequenced tiger trio had a mean depth of 62× (range: 58×–67×), while the snow leopard individuals had a mean depth of 40× (range: 32×– 48×). The jaguar and clouded leopards were sequenced to lower depths: the jaguar genomes had a mean depth of 16× (range: 12×–20×), while the clouded leopard genomes had a mean depth of 22× (range: 16×–32×). We compensated for the lower depths by sequencing multiple trios for these species and adjusting the depth and genotype quality filters (See *Supplementary Information*). Notably, the two *Neofelis* siblings show similar inferred per-generation mutation rates despite the difference in mean depths (Sa Ming: 4.89e-09 against mNeoNeb1; Ta Moon: 7.61e-09; Figure S6). All four jaguar trios show similar inferred rates (4.93–7.29e-09; Figure S6; Table S13). Moreover, the average jaguar (6.62e-09) and clouded leopard rates (6.23e-09) were within the observed Pantherinae range excluding these species.

### Lion genome assembly

We assembled a novel lion genome using PacBio HiFi (40.55× coverage; Table S5) and Dovetail Omni-C (40.64× coverage; Table S5) technology from a single, male lion known as “Luke”. The final assembly resulted in two phased haplotypes in complete chromosomes (Tables S5 and S9, Figure S2). The contig and scaffold N50 of each haplotype was 36.1 and 148.8Mb for haplotype one and 28.9 and 149.4Mb for haplotype two, respectively (Table S9). Both haplotypes had higher BUSCO scores than previously published lion assemblies, as did the ‘mappable’ assembly, which comprised a single haplotype with both sex chromosomes included (Table S9). Notably, the Y chromosome is the most contiguous assembled felid Y chromosome to date, spanning approximately 22Mb (See *Supplementary Information* for details; Figures S3–S5). Additionally, most scaffolds contained evidence of telomere motifs (Figure S2), confirming that the assembly has high continuity and contiguity.

### DNM rate calculations

As is standard practice in mutation-rate estimation from trios (e.g. Besenbacher et al. 2019; Bergeron et al. 2023), we only consider single-forward mutations occurring at loci where both parents are homozygous, but we report the number of putative double-forward and backward mutations and mutation rates including these loci in the *Supplementary Information* (Table S12-S14). After filtering, we retained a mean of 1,006,206,181 callable sites (standard deviation: 221,106,937; median: 1,048,216,390; range: 588,548,939–1,289,424,826) across all *Panthera* and *Neofelis* analyses (Table S11). We identified between 5 and 116 potential *de novo* mutations per trio across all trios and testing conditions. Further filtering of clustered candidate DNMs and overlaps between siblings (when available) reduced this number to between 3 and 82 potential *de novo* mutations across all trios (Table S12). As we show below, much of the variation in number of DNMs is due to the variation in the number of callable sites across species and genome assemblies. WhatsHap pedigree phasing could not identify the *de novo* parental origin of any of the candidate DNMs.

Mean mutation rates ranged from 2.13e-09 to 3.46e-08 mutations per bp per generation (Table S13) across all trios, genome assemblies, and analyses (Figure 3). Considering only the most contiguous genomes for each species, species-averaged mean rates ranged between 4.97e-09 (snow leopard) and 1.43e-08 (leopard) mutations per site per generation. Average Pantherinae mutation rates were approximately normally distributed (Shapiro-Wilk normality test using *shapiro.test* in R 4.3.1, *W* = 0.8278, *p =* 0.103), with a mean of 8.07e-09 (standard error: 1.41e-9; 95% C.I.: 5.30e-9–1.08e-8), a median of 6.53e-09, a standard deviation of 3.46e-09, and minimum of 4.97e-08 in the snow leopard and maximum of 1.43e-08 in the Bergeron et al. leopard trio.

We observed differences in mean rates between analyses that varied only in the employed reference sequence. The tiger trios had an average mean rate of 9.88e-09 against PanTigT.MC.v3, but an average of mean rate of 2.50e-08 against GenTig1.0 (Figure 2). Notably, the confidence intervals for the Bergeron et al. (2023) tiger trio were non-overlapping with the other tiger analyses (Figure S6). For *Neofelis*, the inferred average mean rates and their corresponding confidence intervals differed between the three different assemblies (Figure 2). While the average mean rates and those for the higher coverage *Neofelis* sibling (Sa Ming: 32×) increased with genome fragmentation (Sa Ming mean rates: mNeoNeb1: 4.89e-09; SNNU_Nneo_1: 6.51e-09; SaMing-1434: 8.10e-09), the lower coverage sibling (Ta Moon: 16×) showed the opposite pattern (Ta Moon mean rates: mNeoNeb1: 7.61e-09; SNNU_Nneo_1: 7.18e-09; SaMing-1434: 6.78e-09). Nevertheless, there was significant overlap between the binomial confidence intervals for the three assemblies, supporting the accuracy of the inferred *Neofelis* mutation rates (Figure 2). Furthermore, we observed that the mutational spectra varied between analyses (Table S14), even when the overall estimated mutation rate was similar, such as in the lions (6.43e-09 average mean rate [range: 5.81e-09–7.05e-09] against the novel Luke assembly versus 7.13e-09 mean rate [range: 5.62e-09–8.64e-09] against PanLeo1.0).

**Figure 2:**
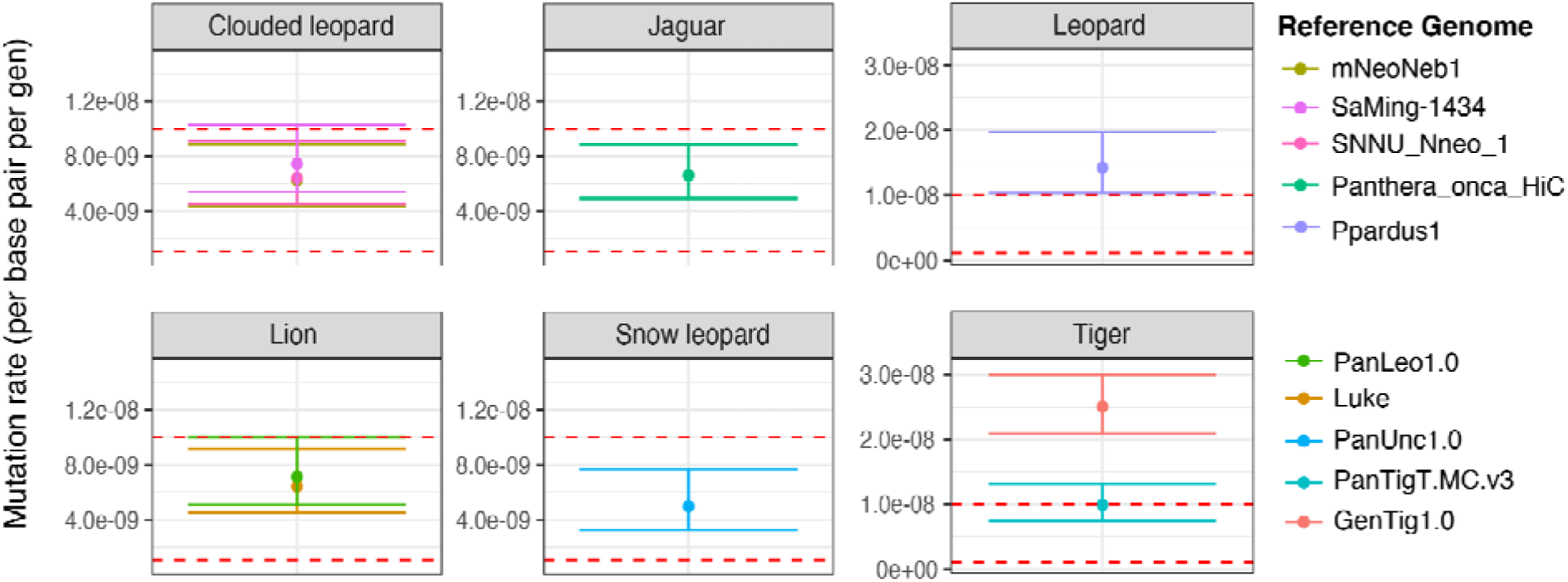
Final mutation rates across all individuals with 95% CIs for each species colored by genome assembly used. Red dotted lines represent the previously inferred mutation rate minimum (1.1e-09) and maximum (1.0e-08) for the Pantherinae lineage.

**Figure 3:**
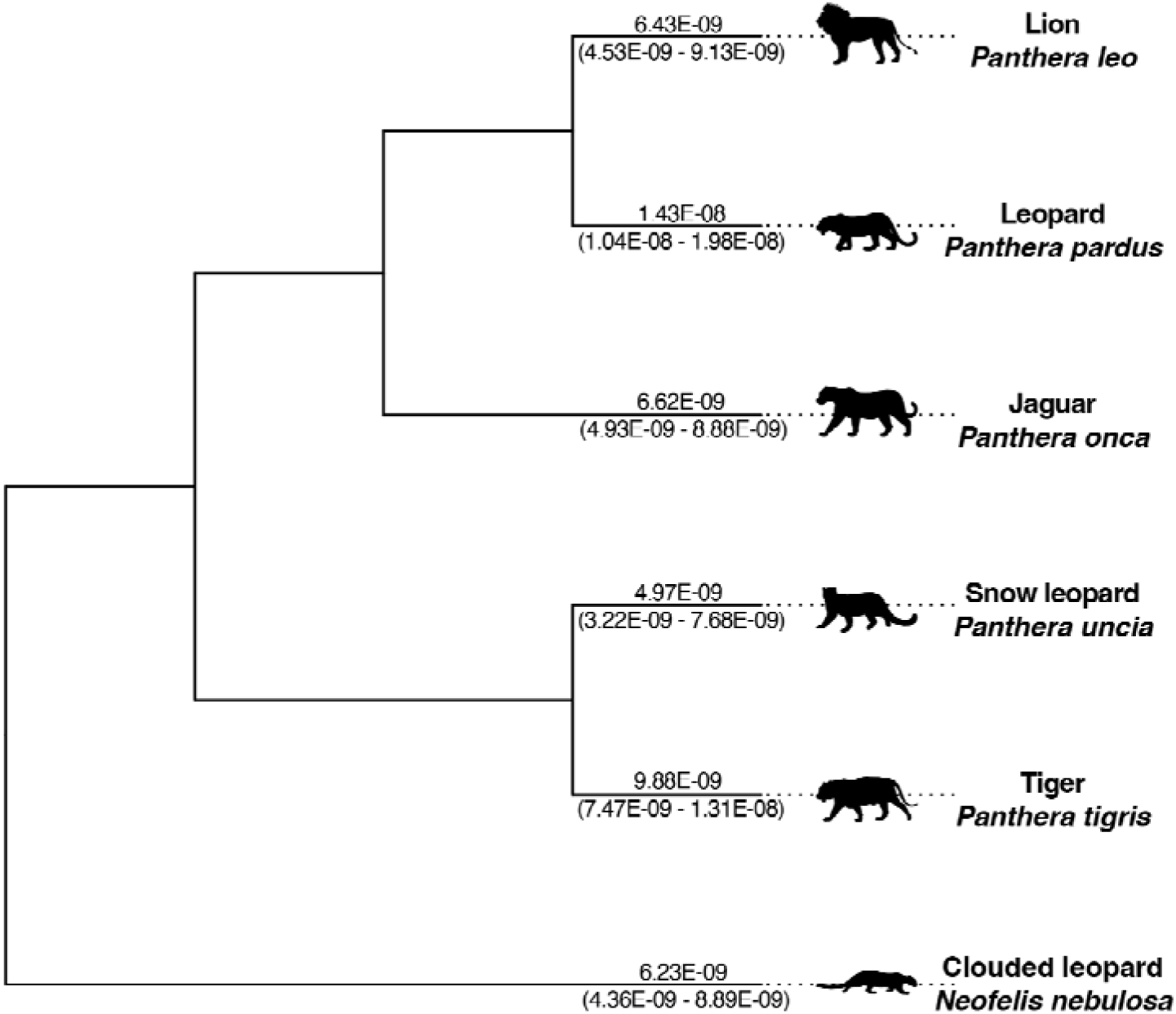
Mean inferred germline mutation rates (per bp per generation) across species overlaid on a dendrogram of the Pantherinae. Lineage-specific rates are reported as averages from the highest quality genome assembly across individuals for that species. Silhouettes were obtained from PhyloPic under a CC BY-NC 3.0 DEED license. Specific image attributions can be found in the acknowledgements.

### Demographic history

We used PSMC to investigate historic effective population size (N_e_) changes across the Pantherinae contingent upon the applied mutation rate estimated for each species. Coalescent methods scale the inferred N_e_ after estimating diversity (θ) in windows across the genome. The method then scales the results to years and effective population size using the germline mutation rate (µ) and generation time through the equation θ = 4N_e_µ. Previously used mutation rates resulted in nearly an order of magnitude difference in the historic effective population size predictions (Figure 1), which is unsurprising given the differences in previous mutation rate inference (Table S1). Mutation rate differences also shift the timing of the demographic events, where slower rates push back the approximate timing of events. Each species’ inferred demographic history reveals population declines occurring approximately 100,000 and 10,000 years ago (Figure 4B, Figures S7–S9), coinciding with megafauna declines which have been attributed to human migrations out of Africa and/or the last glacial maximum (Bergman et al. 2023). The snow leopard has the lowest predicted past effective population size of any of the species examined as previously reported (Solari et al. 2023), while the African leopard and lion have the highest predicted past population sizes.

**Figure 4:**
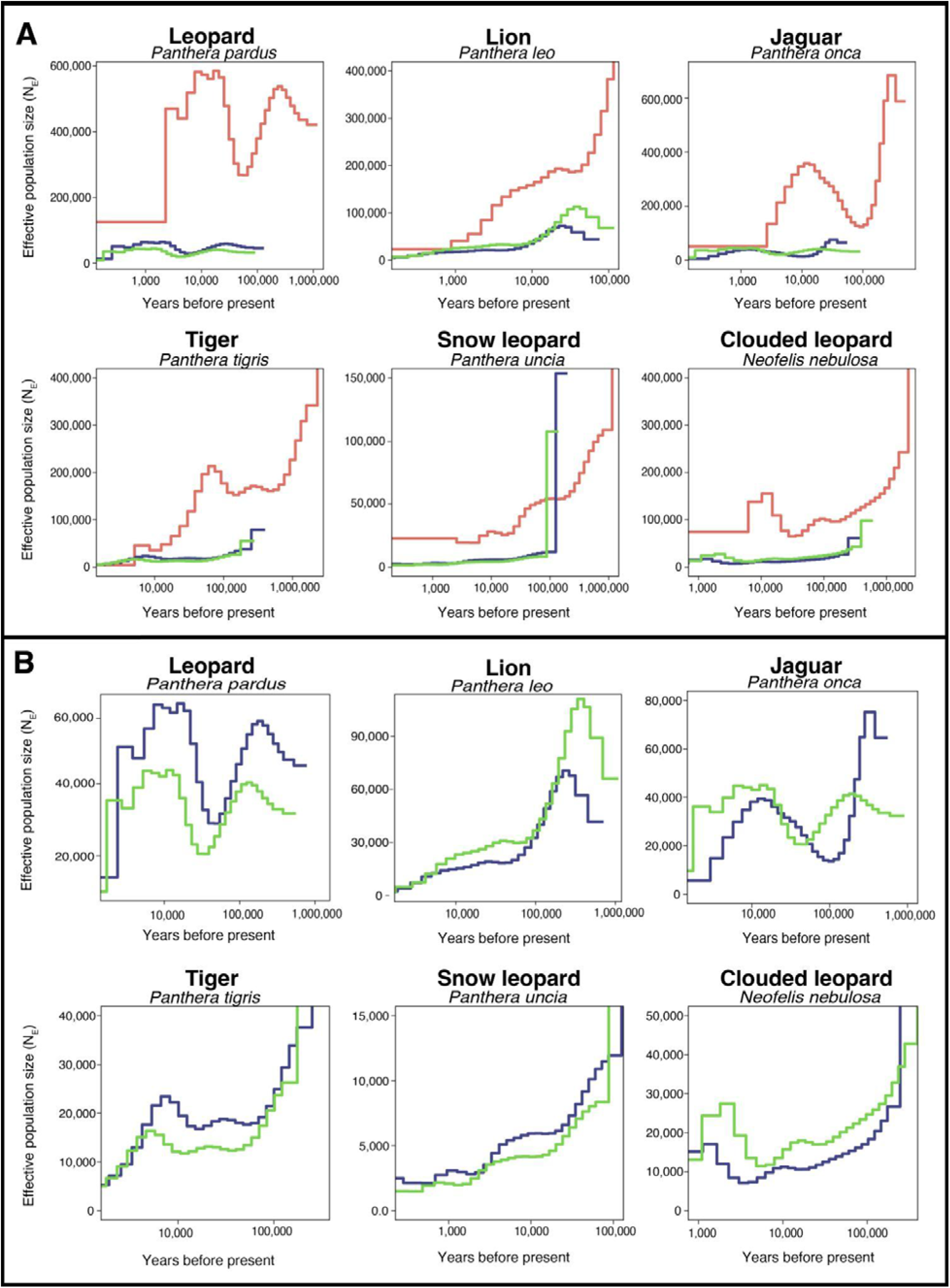
Plots of inferred historic population sizes for individual species of Pantherinae reconstructed with PSMC. Note that the x and y axes are different for different species. Maximum mutation rate (1.0e-08 per base pair per generation) is shown in dark blue, minimum rate (1.1e-09 per base pair per generation) is shown in pink, and the newly inferred average rate (Figure 3) is shown in green. Panel A shows all rates, Panel B shows only the maximum rate and new rate so that the differences can be more easily observed.

## Discussion

Germline mutation rates are a crucial parameter for inferring divergence times and predicting past population sizes (Li and Durbin 2011; Terhorst et al. 2017; Schiffels and Wang 2020) as well as the spectrum of mutagenesis more generally (Carlson et al. 2020). Using whole-genome pedigree data representing six of the seven species of Pantherinae (the missing species being the Sunda clouded leopard, *Neofelis diardi*), we found that *de novo* mutation rates are broadly similar across the clade, with the notable exception of the leopard’s being somewhat higher. However, the parents in the leopard trio (mean age: 9.425 years) were much older than the mean ages of the parents in the other trios (lion, 5.5; tiger, 5; snow leopard, 4; jaguar, 5.8; clouded leopard, 2.7; Table S4). Mutation rates are known to increase with parental age (e.g. Bergeron et al. 2023), so age may be the cause of this apparent increase rather than lineage rate acceleration. Additionally, we show that multiple offspring and/or distinct pedigrees are useful for reducing false positive mutations, neither of which were available for the leopard. The low level of variation in mutation rates between Pantherinae species is consistent with Bergeron et al. (2023), who found that terrestrial mammal mutation rates are relatively constrained compared to other amniotes.

Our inferred per-generation mutation rates for the published tiger and leopard trios are higher than those reported by (Bergeron et al. 2023). This can largely be attributed to pipeline differences in determining the number of callable sites. The total number of inferred DNM candidates were very similar between the two studies (tiger: 29 candidates in Bergeron et al. vs. 24 identified using RatesTools; leopard: 35 in Bergeron et al. vs. 37 using RatesTools). However, Bergeron et al. did not remove low-mappability (i.e. regions which are not unique enough to reliably map reads) and repetitive regions, resulting in much higher estimates of the callable genome (tiger: 90% callable in Bergeron et al. vs. 52% using RatesTools; leopard: 80% in Bergeron et al. vs. 53% using RatesTools). Prior to filtering these low-mappability and repetitive regions, RatesTools retains more similar proportions of the genome to Bergeron et al. (tiger: 92%; leopard: 94%). It has been established previously that repetitive regions in the genome have poor mapping quality compared to other regions and cause errors in variant calling (Krusche et al. 2019) even at very high (>30×) coverages. It is therefore likely that inclusion of these regions could lead to unpredictable bias in the rate estimate. The regions of the genome that should be excluded for high-confidence variant calls is debated on even for systems with telomere-to-telomere genome assemblies and many high-coverage individuals (https://www.illumina.com/science/genomics-research/articles/identifying-genomic-regions-with-high-quality-single-nucleotide-.html). Given these considerations, we suggest our conservative approach to region inclusion likely reduces these potential biases, especially in the absence of PCR verification. However, additional studies utilizing high-coverage and multiple long-read sequencing technologies, in addition to PCR validation of candidate *de novo* mutations, would provide a much-needed gold-standard for the field (e.g., Noyes et al. 2022).

We observed differences in inferred mutation rates across different genome assemblies. In the tiger and clouded leopard trios, the lower-quality assemblies (GenTig1.0 and SaMing- 1434) had higher inferred mean mutation rates compared to the higher-contiguity assemblies (Table S13), though they have substantial overlap in their confidence intervals, except in the case of the tiger data from Bergeron et al. 2023 (Figure 4). In the clouded leopard analysis, we inferred slightly differing rates even between the two long-read assemblies. While the inferred mutation rates between the two lion assemblies were similar, the mutational spectra differed (Table S14), showing that the reference assembly impacts *de novo* mutation identification even when not changing the overall mutation rate. Nevertheless, we argue that our inferred tiger mutation rates against PanTigT.MC.v3 are accurate given that: (1) they are highly consistent between the two completely independent trios (10.3e-9 and 9.52e-09 for the novel and Bergeron trios respectively), (2) the rates are similar to those determined via the pipeline used in Bergeron et al. taking into account the differences in number of callable sites (see above), and (3) they fall within the range of the other Pantherinae rates reported here. Similarly, while we observed some differences in the inferred rates and mutational spectra for clouded leopards and lions, the average rates and confidence intervals between the different assemblies are close to each other and fall within the typical Pantherinae range. Encouragingly, the inferred trio-based lion mutation rate (6.43e-09) is very similar to that estimated using neutral divergence (5e-9) by (Cho et al. 2013).

While we do not have direct evidence that reference bias (in which the quality or identity of the reference influences the ability to accurately detect variants) could impact inferred *de novo* mutation rates, our tiger and clouded leopard results suggest that this may be the case, as have previous studies (Wang et al. 2021). We speculate that for species with strongly divergent subspecies and populations, alignment of a trio to a reference genome built from a different subspecies/population may result in increased genotyping error and impact downstream mutation rate estimates (Thorburn et al. 2023). Notably, the tiger trio from (Bergeron et al. 2023) was more impacted by alignment to the GenTig1.0 generic tiger reference (3.6-fold rate increase over alignment to PanTigT.MC.v3) than was our newly reported tiger trio (1.4-fold increase). GenTig1.0 was assembled from Kylo-Ren, the offspring included in our trio. Similarly, the clouded leopard trios had different inferred rates between the mNeoNeb1 and SNNU_Nneo_1 assemblies, despite overall similar genomic assembly qualities. This may be attributable to the SNNU_Nneo_1 assembly being a different *N. nebulosa* population (from China) (Yuan et al. 2023), while the mNeoNeb1 assembly was built from the same population (from Thailand). The increasing mutation rate with genome fragmentation in the higher coverage sibling is likely due to an increased rate of misalignment and inaccurately called SNPs, while the decreasing rate in the lower coverage sibling is likely due to an increased rate of missing heterozygotes (e.g. Maruki and Lynch 2017; Rhie et al. 2021). Overall, our results make it apparent that multiple trios aligned to high-quality genomes are preferable for reducing the error associated with direct estimates of mutation rate. High quality genomes with respect to contiguity and continuity and high-coverage data allow for the inclusion of more callable sites (Table S11) and reduce the chance that heterozygotes are missed or miscalled. We encourage future trio-based *de novo* mutation analyses to test multiple genome assemblies to help determine whether rate estimates are robust to reference genome choice.

Given the rarity of *de novo* mutations, analyses of single trios can result in extreme values simply by chance. For instance, the leopard mutation rate is an outlier for the Pantherinae, but we cannot currently determine whether this is a chance event, an age effect, or rate acceleration in *Panthera pardus*. We therefore strongly encourage the sequencing of multiple trios per lineage to better infer rate variation. In many cases, a single sequenced trio per species may be all that is available (Bergeron et al. 2023; Prado et al. 2023). In these cases, an averaged rate across multiple closely related species may be preferable to avoid inferred extreme rates arising simply by chance. Furthermore, we recommend that researchers carefully consider whether published mutation rates from trios are appropriate for their applications. Given that inferred mutation rates depend both on the filtering parameters and reference genome choice, researchers may need to recalculate the rate using the published trio raw data (e.g. if they are using a new genome assembly or very different filtering parameters) or adjust their analysis parameters to match the assumptions of the published rate.

One limitation of our analyses is we were unable to determine the parental origin of *de novo* mutations despite pedigree phasing using WhatsHap. A paternal bias in mammalian germline mutations has been observed in a wide variety of amniote species (e.g. de Manuel et al. 2022). Future trio analyses using long-read sequencing may help determine whether such a bias is also observed in Pantherinae. Phasing could also be improved by the analysis of a greater number of individuals in the pedigrees.

Finally, we explored the impact of mutation rates in the context of a commonly used historic demographic inference tool, PSMC. While we acknowledge that other methods now exist, including several varieties of PSMC (Cahill et al. 2016; Liu et al. 2022; Cousins et al. 2024) and those which use population-level data (Browning and Browning 2015; Terhorst et al. 2017; Schiffels and Wang 2020), we chose to use PSMC to illustrate this issue because it remains a commonly used tool, and one of few tools appropriate for inferring historic effective population size from single genomes. We show that, not unexpectedly, the order of magnitude rate difference of previously published mutation rates substantially impact the inference of demographic history (Figure 4) by changing the inferred effective population size and the timing of demographic events, which has been previously observed (Nadachowska-Brzyska et al. 2016). These new rates will provide a solid foundation for further exploration of the demographic history in the Pantherinae.

In conclusion, our results narrow the range of the previously inferred mutation rate for the Pantherinae (between 0.5 and 1.4e-08 per base pair per generation, mean 0.81e-08 ± 0.35- 08). We provide insight into the possible caveats of direct mutation rate estimation and its impact on historic demographic inference. Notably, we find that the choice of reference genome and depth of coverage has a substantial impact on the ability to detect *de novo* mutation candidates. This complicates cross-species rate comparisons as it is difficult to precisely control genome assembly quality between organisms with differing genomic architectures and base compositions. Furthermore, the availability of high-quality DNA samples for chromosomal-level assembly varies widely across study systems, producing inherent biases in genome assembly qualities. We therefore urge researchers to assess available genome assembly quality through careful curation (Howe et al. 2021), the inclusion of multiple parent-offspring trios, and the quality and depth of the trio sequencing data before inferring inter-species mutation rate change. Careful consideration of these factors will provide credence to future demographic models of non-model organisms.

## Acknowledgments

The Woodtiger Fund, the Smithsonian Institution, and the following grants to DAP: NIH/NIGMS grant no. R35GM11816506, the Chan Zuckerberg Initiative, and National Science Foundation (NSF) grant no. 2225088. E.E.A. was supported by the NSF GRFP and a Washington Research Foundation Postdoctoral Fellowship. Additional support for analysis was provided by the United States Department of Agriculture National Institute of Food and Agriculture Postdoctoral Fellowship no. 2022-67012-38987 (S.B.C.), NSF IOS-PGRP CAREER no. 2239530 (A.H.), and CNPq/Brazil grant no. 309068/2019-3 (EE). Mutation rate computations were performed on the Smithsonian Institution High-Performance Computing Cluster (doi: 10.25572/SIHPC). Additional computing for this project was performed on the Sherlock cluster. We would like to thank Stanford University and the Stanford Research Computing Center for providing computational resources and support that contributed to these research results. We thank the Smithsonian’s National Zoo and Conservation Biology Institute for providing the lion and mainland clouded leopard samples. We are specifically grateful for the assistance of Adrienne Crosier, Jennifer Donato, Erin Latimer, Kelly Helmick, Mandy Murphy, Craig Saffoe, and the staff of the NZCBI Department of Wildlife Health Sciences in obtaining the NZCBI samples and their associated metadata. We are indebted to Tigers in America and In-Sync Exotics (Wylie, Texas) for samples from Kylo-Ren, Zahra, and Assad. We are very grateful to the San Francisco Zoo for providing samples for Anna, Raj, Tikka, and Tsering. We thank Sebastián di Martino, Emiliano Donadio, Fundación Rewilding Argentina, Henrique Figueiró, Caroline Sartor and Giovanna Oliveira for access to jaguar samples and their laboratory processing. We also thank Elizabeth Hadly, Molly Schumer, and Noah Rosenberg for helpful conversations and advice. Whole-genome resequencing of the clouded leopard individuals was supported by the Biodiversity Genomics Exemplar Project and the Smithsonian Institution Grand Challenges Award Program. We thank Erich Jarvis, Olivier Fedrigo, and the staff of the Vertebrate Genome Laboratory at Rockefeller University, New York, for the sequencing, assembly, and curation of the clouded leopard mNeoNeb1.pri assembly. PhyloPic image credit to: *Panthera onca* and *Panthera uncia*, Gabriela Palomo-Munoz; *Panthera pardus*, Lukasiniho; *Panthera leo*, *Panthera tigris*, and *Neofelis nebulosa* were obtained under a CC0 1.0 DEED public license.

## Author contributions

E.E.A. and M.G.C. wrote the paper and prepared the figures and tables. E.E.A. and M.G.C. wrote and maintain the RatesTools pipeline. E.E.A., S.B.C., and A.H. assembled the newly reported lion genome. E.E.A. and M.G.C. performed the bioinformatic analyses. E.E.A., K.A.S., K.-P.K., M.G.C., and G.Z.L. performed the genomic experiments. N.A. and P.W. performed the autopsy of the lion “Luke”. R.C.F., J.E.M., N.A., P.W., K.A.S., K.-P.K., G.Z.L., and E.E. provided samples and genomic sequences. E.E.A., D.A.P., R.C.F., J.E.M., E.E., K.-P.K., and M.G.C securing funding for this research. D.A.P. and M.G.C. supervised the project. All authors revised and approved the final manuscript.

## Data availability

Newly generated sequence reads are available under BioProject PRJNA1092733. The newly reported lion assembly is available under BioProjects PRJNA1095652 (haplotype 1 with Y chromosome) and PRJNA1095651 (haplotype 2 with X chromosome). The list of lion y-mers, mappable novel lion assembly, and RatesTools configuration files can be found in the Smithsonian Figshare repository under DOI: 10.25573/data.25374244.

## Notes

### Competing Interest Statement

The authors have declared no competing interest.

