## Supplemental Information for "Parameterizing Pantherinae: *de novo* mutation rate estimates from *Panthera* and *Neofelis* pedigrees"

Supplementary Information Files for Panthera Mutation Rates

### **Table of Contents**

1. RatesTools Updates
2. RatesTools Analyses Partitioned by Library Type
3. Minimum Depth and Genotype Qualities for Low-Coverage Trios
4. Supplementary Tables and Figures

### **1. RatesTools Updates**

Since the initial publication of the RatesTools pipeline described in Armstrong & Campana 2023, we have significantly revised the pipeline to streamline code, correct software bugs, and add new features. Complete documentation of the code revisions are available in the GitHub repository (<https://github.com/campanam/RatesTools>). As of version 1.0.0, RatesTools has been upgraded to use Nextflow (Di Tommaso et al. 2017) domain-specific language (DSL) 2. Default software versions for dependencies have been updated to more recent releases: SAMtools, BCFtools and HTSlib 1.18 (Danecek et al. 2021), Genome Analysis Toolkit 4.4.0.0 (McKenna et al. 2010), Picard 3.1.0.7 (Broad Institute 2023 <https://broadinstitute.github.io/picard/>), Java 1.17, Sambamba 0.8.2 (Tarasov et al. 2015), RepeatMasker 4.1.5 (Smit et al. 2013-2015), RepeatModeler 2.0.5 (Flynn et al. 2020), Nextflow 23.10.0, and Ruby ([www.ruby-lang.org](http://www.ruby-lang.org)) 3.2.2.

The initial alignment processes have been streamlined into fewer piped steps to improve runtimes and reduce disk-space footprints. After chromosome filtration, there is now an optional step to perform pedigree phasing of the autosomal variant call format (VCF) file using WhatsHap 2.1 (Garg et al. 2016; Martin et al. 2016). After initial DNM candidate identification, the dnm_summary_stats.rb script provides further identification of classes of mutations for double-forward and backward mutations. Optionally, the dnm_summary_stats.rb identifies and removes clumps of candidate *de novo* mutations using a user-specified distance (in base pairs). These likely represent misalignments or complex de novo mutations rather than multiple single-nucleotide mutations. The script will also identify and remove shared DNM candidates between siblings as these likely represent a genotyping error in one of the parents. After removal of these likely errors, the script will adjust the point-estimate, the bootstrapped mean estimate of the mutation rate, and standard error by multiplying the initial estimates by the fraction of candidates that remain after these filters. An adjusted bootstrapped 95% confidence interval is estimated by multiplying the adjusted standard error by 1.96 and adding and subtracting this critical value from the adjusted bootstrap mean. Additionally, binomial confidence intervals for the mutation rate estimates are calculated using the *binconf* function in the R package Hmisc 5.1-1 (Harrell et al. 2018). Furthermore, the dnm_summary_stats.rb script now calculates mutational spectra for *de novo* single-nucleotide polymorphism candidates.

### **2. RatesTools Analyses Partitioned by Library Type**

For the newly reported lion and tiger trios, we generated both 10x Genomics Chromium and standard Illumina libraries for all three tigers (Assad, Zahra, and Kylo-Ren) and the lion parents (Luke and Nababiep). As an internal control, we ran RatesTools using the parameters described in the primary text, separating the datasets by library type (with the exception that the standard Illumina lion offspring data were used in both partitioned lion analyses). As expected, we retained less of the callable genome in the partitioned datasets than in the combined analyses (25%–44% compared to 45%–51%: Table S11) as the minimum depth (20× per individual) was not adjusted for the lower coverage of the partitioned analyses. A naïve expectation would be an increased observed mutation rate in the lower-coverage datasets due to increased impact of genotyping errors (Maruki and Lynch 2017). Counter-intuitively, the observed mean mutation rates decreased in each of the partitioned analyses compared to the complete datasets, likely because the decreased coverage increased the false negative rate in most analyses due to our stringent depth and genotype quality filters (Tables S3 and S13). For instance, the Koch et al. (2019) DNp statistic filters candidate DNMs based on relative likelihoods of the possible genotypes, so it is less likely to retain a candidate DNM with lower-depth and quality (Koch et al. 2019). Nevertheless, the binomial confidence intervals of the partitioned analyses always encompassed the calculated mean rate for the unpartitioned analyses, suggesting that our overall rate estimates are robust.

Furthermore, we observed shared candidate mutations between these pseudo-replicate analyses. Against the novel Luke lion genome assembly, we observed 18 candidate DNMs in the 10x Genomics Chromium data and 8 in the standard Illumina data. Of these, 3 were detected in both datasets. Against the PanLeo1.0 assembly, we identified 13 candidate DNMs each in the 10x and standard Illumina data, of which 2 were shared. For the novel tiger trio aligned against the PanTigT.MC.v3 assembly, we identify 10 candidate DNMs in the 10x data and 14 in the standard Illumina data, of which 2 were shared. When aligned against the GenTig1.0 assembly, we observe 7 tiger candidate DNMs in the 10x data and 26 in the standard Illumina data, of which 3 were shared. While this is a low rate of replication, it also likely reflects a high false-negative rate in the lower-coverage partitioned datasets.

### **3. Minimum Depth and Genotype Qualities for Low-Coverage Trios**

Variant filtration parameters have a significant impact on inferred *de novo* mutation rates (e.g. Bergeron et al. 2022). Appropriate optimization of these parameters is non-trivial. For our higher-depth trios (lions, tigers, and snow leopards), we chose minimum depth (20×) and genotype quality (GQ: 65) cut-offs that are broadly suitable for high-quality data and similar to those in the published literature (e.g. Wang et al. 2021; Bergeron et al. 2023). These parameters were not suitable for our jaguar and clouded leopard trios which were sequenced to lower depth. We chose a minimum depth of 10× as this is approximately half the depth of most of the lower-coverage genomes (Bergeron et al. 2023) and is a minimum depth for reasonable genotype quality (e.g. Maruki and Lynch 2017). The GQ is dependent on the depth of sequencing (McKenna et al. 2010; Caetano-Anolles 2023 <https://gatk.broadinstitute.org/hc/en-us/articles/360035890451-Calculation-of-PL-and-GQ-by-HaplotypeCaller-and-GenotypeGVCFs>; Wang et al. 2021), so it also must be adjusted for sequencing depth as a minimum GQ of 65 is inappropriate for lower-depth sequencing (mean across individuals less than 20×) and produces many false negatives. Conversely, we observed that in genomes above 20× depth, the median GQ exceeded 65 in all cases (Table S3), with most higher-coverage individuals having median GQ values of 99 (the maximum calculated by GATK: Table S3) This means that a lower GQ cut-off (e.g. 40) greatly increases the false positive rate in these genomes as the GQ value is inflated by the higher sequencing depth. We empirically chose a minimum GQ of 45 for the low-depth trios based on the observed mean, median, and first quartile GQ values in our datasets. This value also empirically provided a reasonable balance between false negatives and positives.

### **4. Supplementary Tables and Figures**

**Table S1**: Previously published estimated mutation rates of the genus *Panthera* and methods used.

**Table S2:** Sampling details for newly sequenced trios used in this study

**Table S3**: Genome depth and genotype quality statistics across individuals with data generated in this study.

**Table S4**: Paternal and maternal ages across all trios.

**Table S5**: PacBio and OmniC data statistics.

**Table S6**: Sample details for Y-mer discovery

**Table S7**: Coverage of Y-mers per contig

**Table S8:** List of Y-hinted contigs that could not be incorporated into the Y during scaffolding

**Table S9**: Final assembly statistics for the completed lion genome compared with other publicly available lion assemblies.

**Table S10**: Final assembly BUSCO scores for the completed lion genome compared with other publicly available lion assemblies.

**Table S11:** Number of retained sites at each stage or the RatesTools pipeline.

**Table S12:** Total numbers of DNM sites before and after removing clustered candidates and those shared between siblings.

**Table S13:** Final estimated per-trio mutation rates.

**Table S14:** Mutational spectra of observed candidate *de novo* mutations after filtering sibling-shared and clustered mutations.

**Table S15:** Comparison of 10x Chromium retained callable genome sites with and without barcode trimming.

**Table S16**: Details for individuals used in PSMC analyses

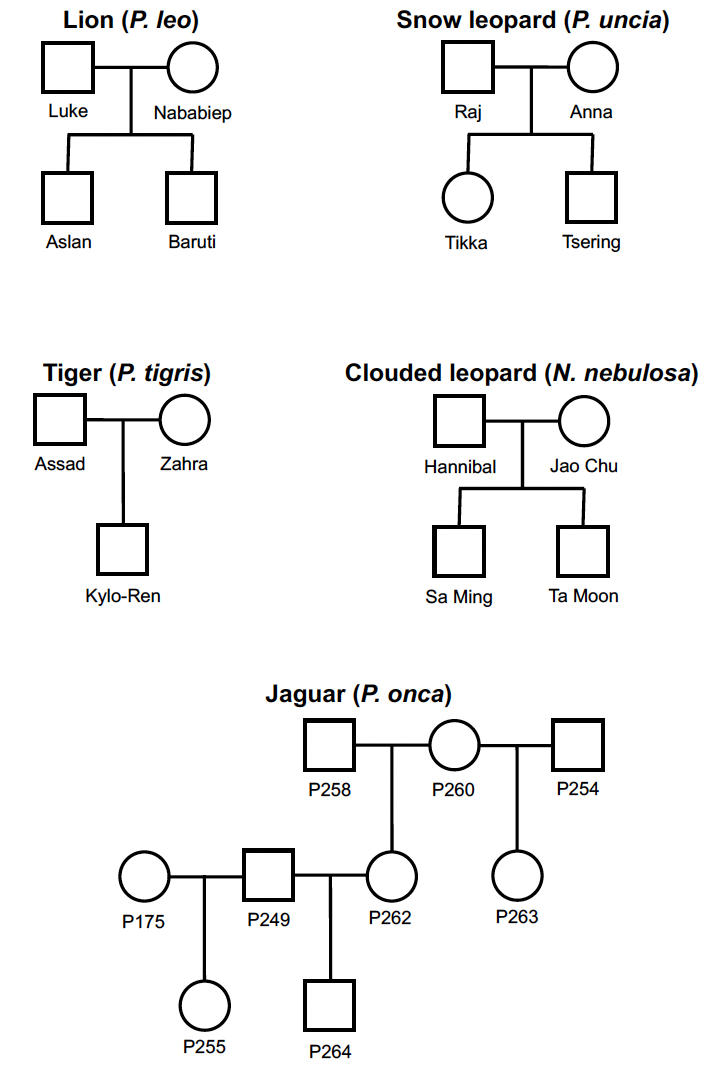

**Figure S1**: Pedigrees of newly reported *Panthera* and *Neofelis* trios.

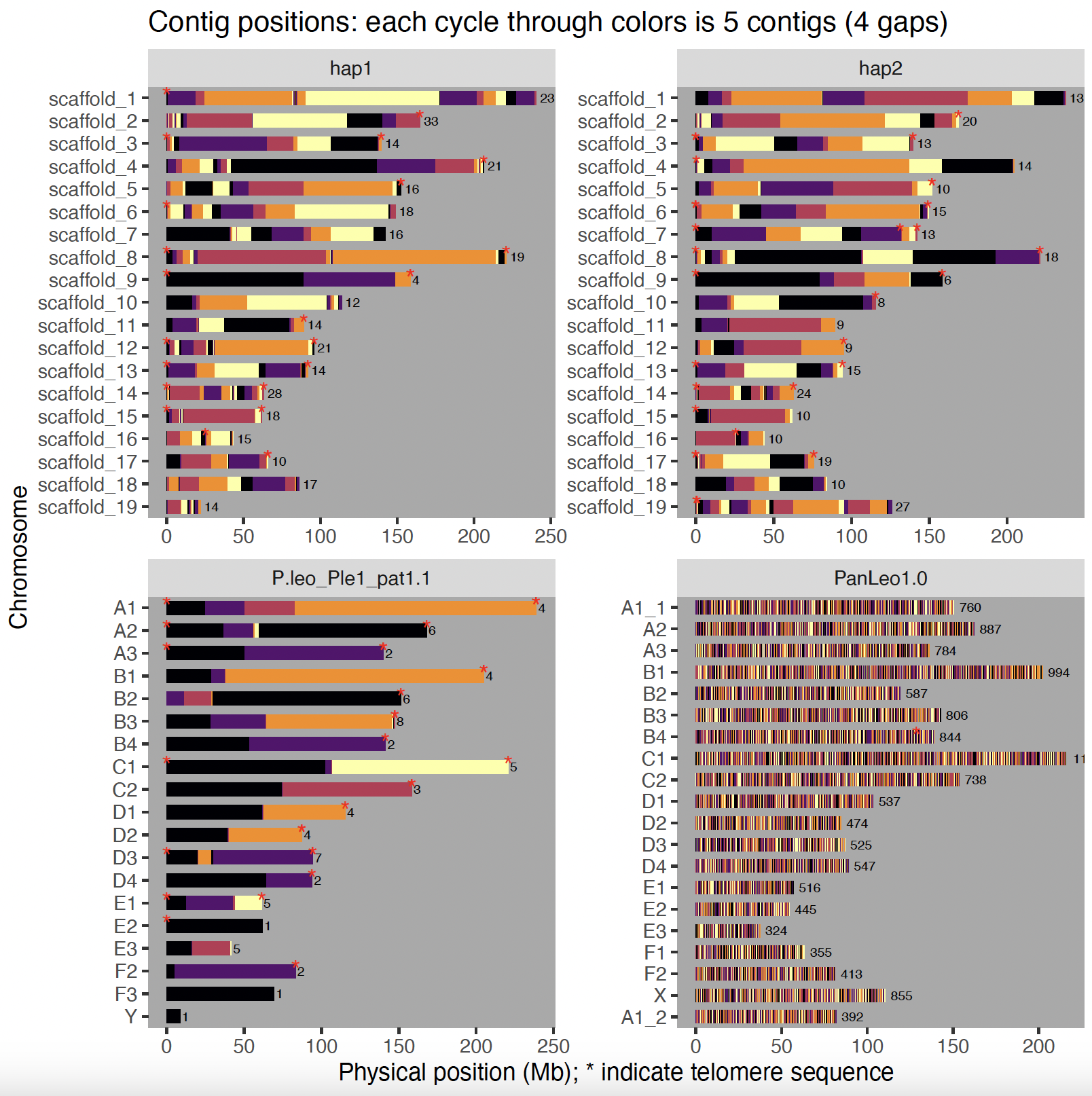

**Figure S2**: Panels showing contig positions across various lion genome assemblies. Top panels represent haplotypes of the newly assembled genome in this study. Red stars represent evidence of telomere motifs.

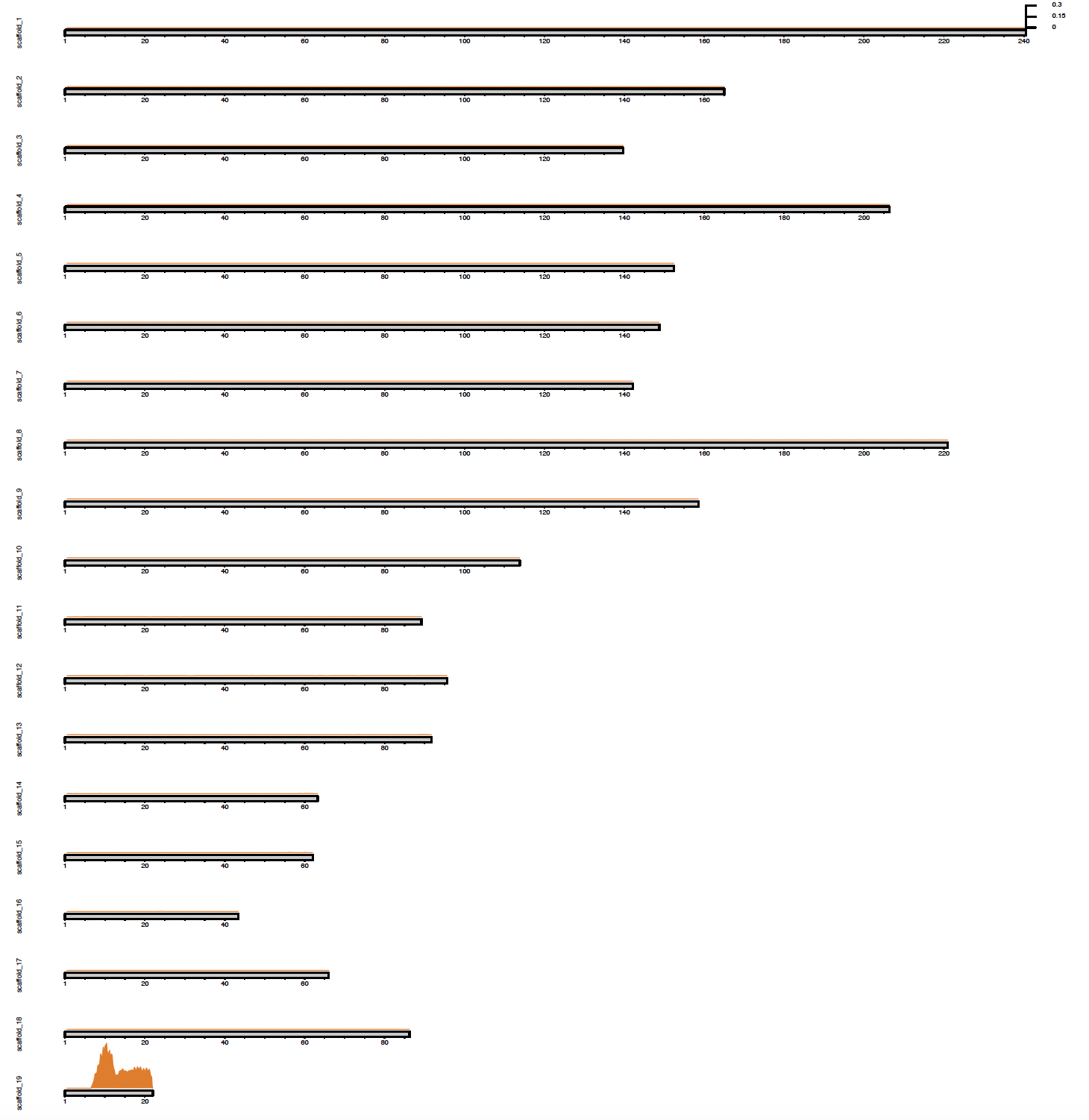

**Figure S3:** Y-mer density plot for haplotype 1.

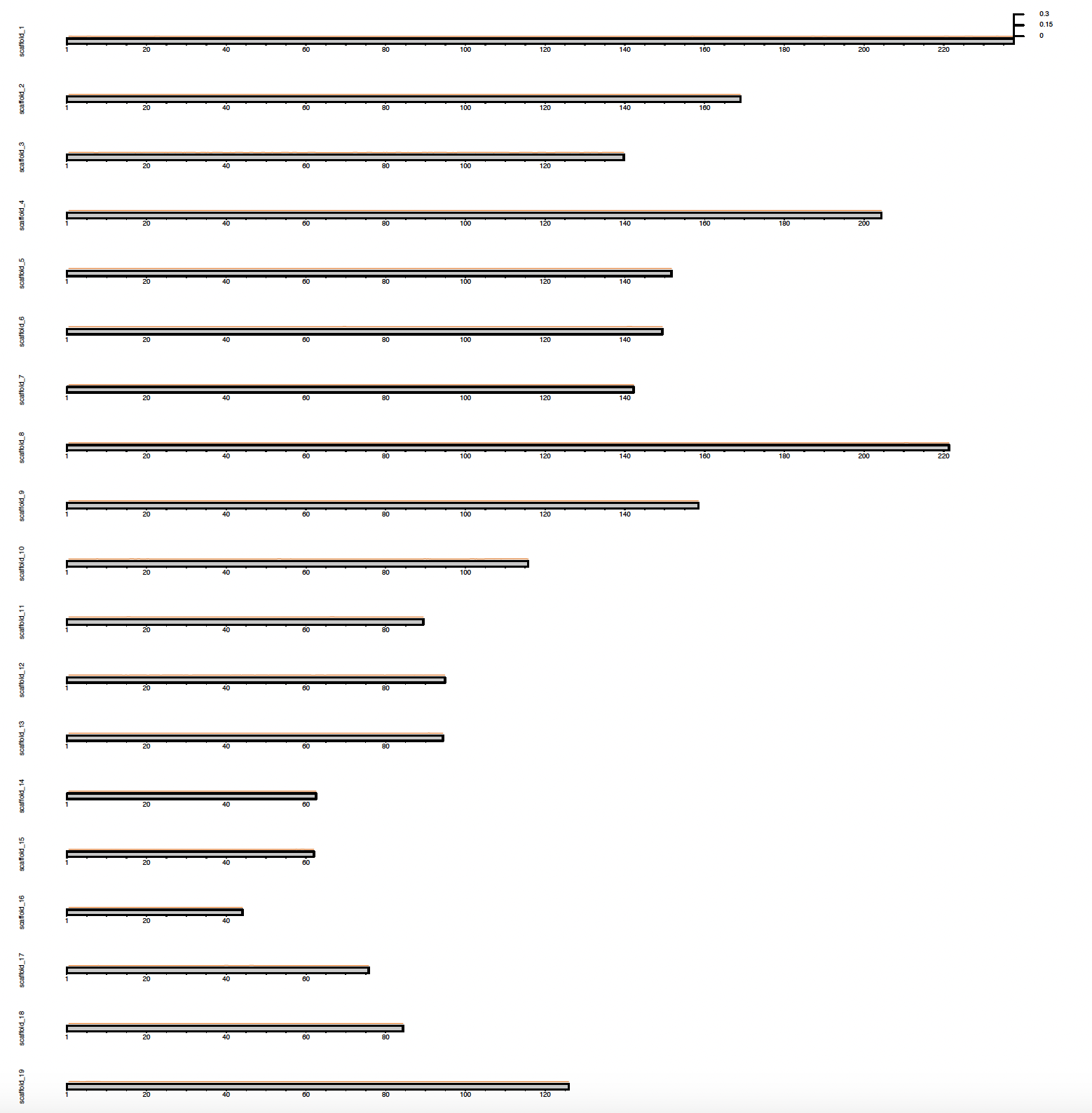

**Figure S4**: Y-mer density plot for haplotype 2.

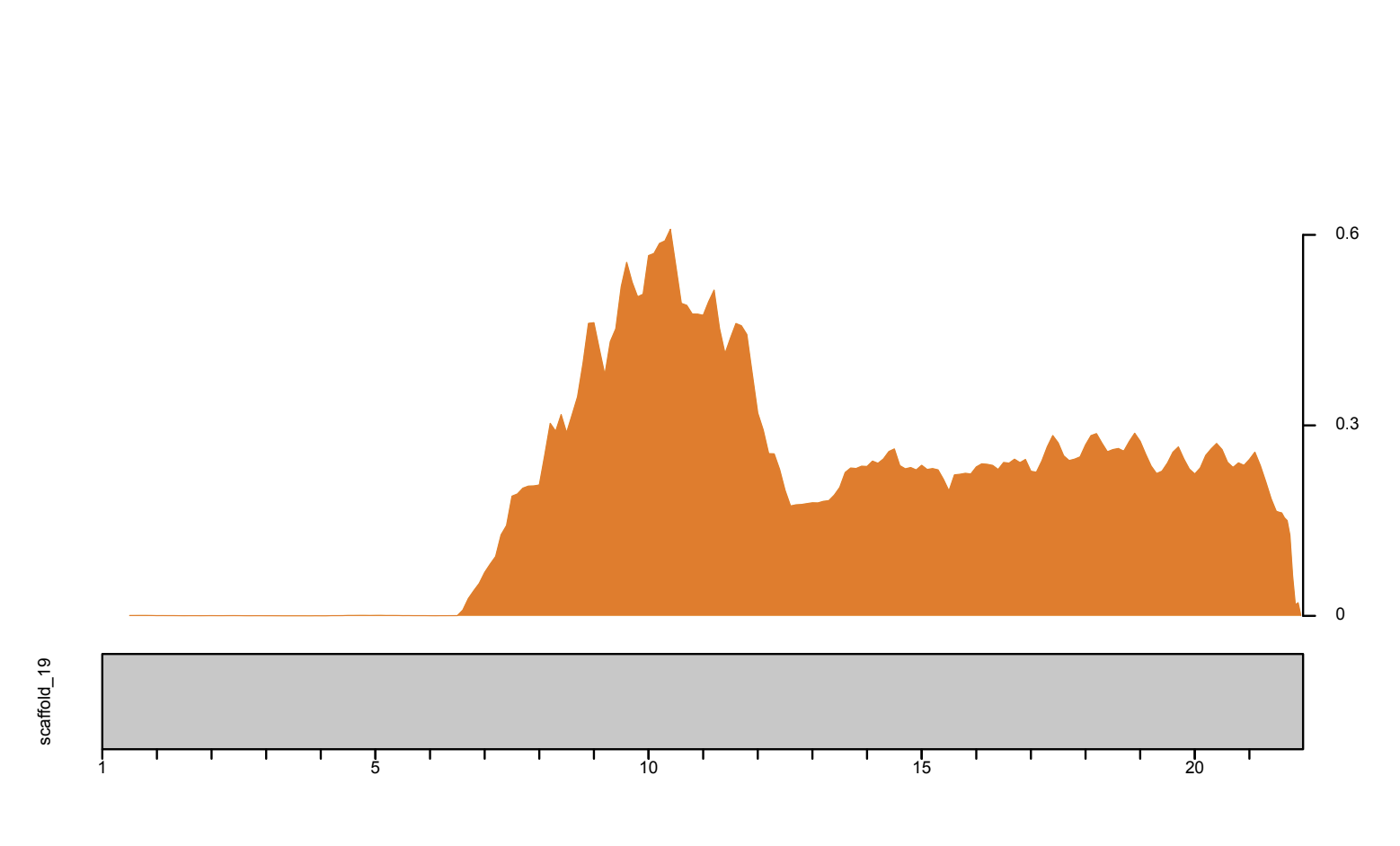

**Figure S5**: Y-mer density plot for Y chromosome. Note that the region without y-mer density represents the pseudoautosomal region (PAR).

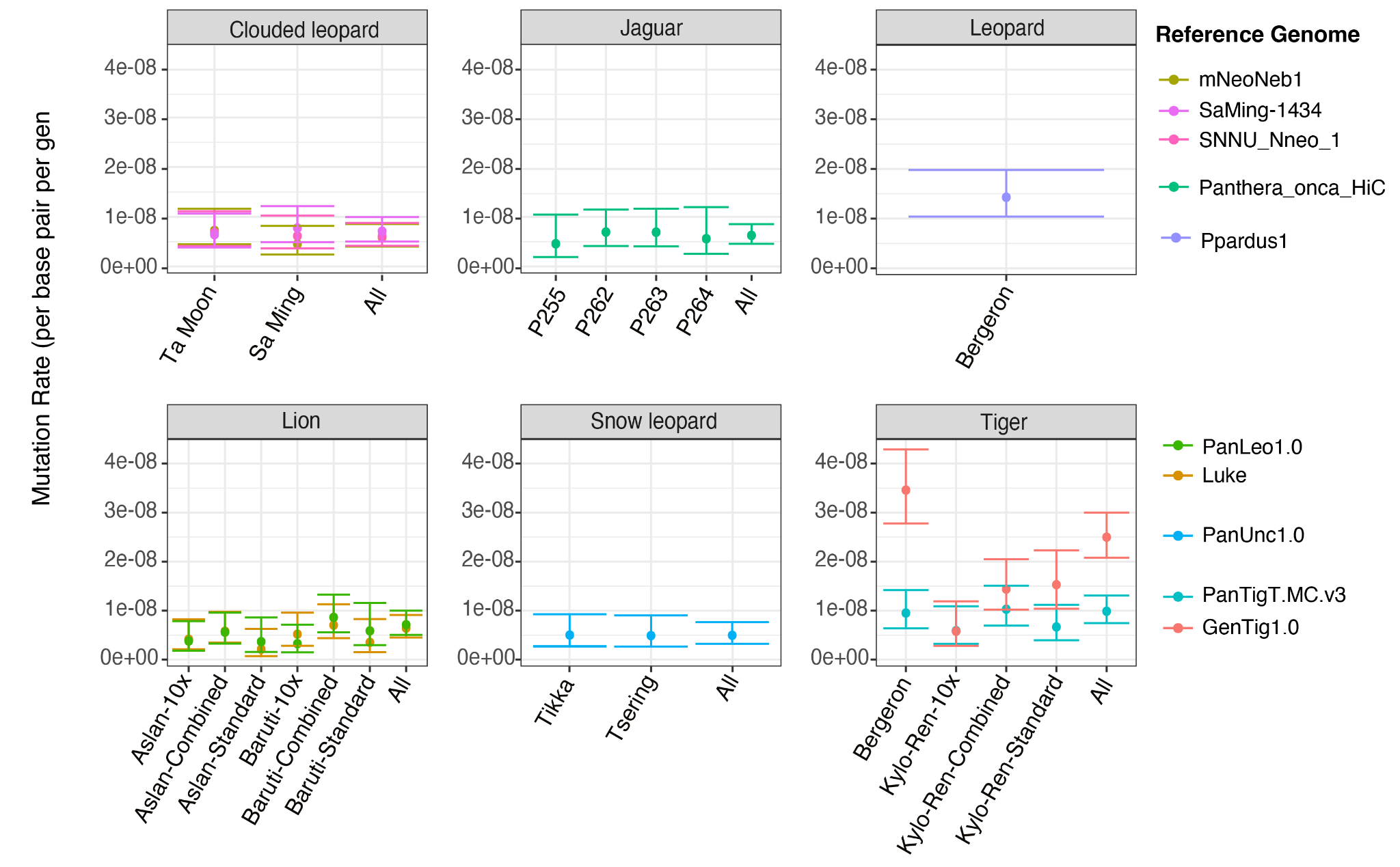

**Figure S6**: Variance across data type and genome assembly on inferred mutation rates.

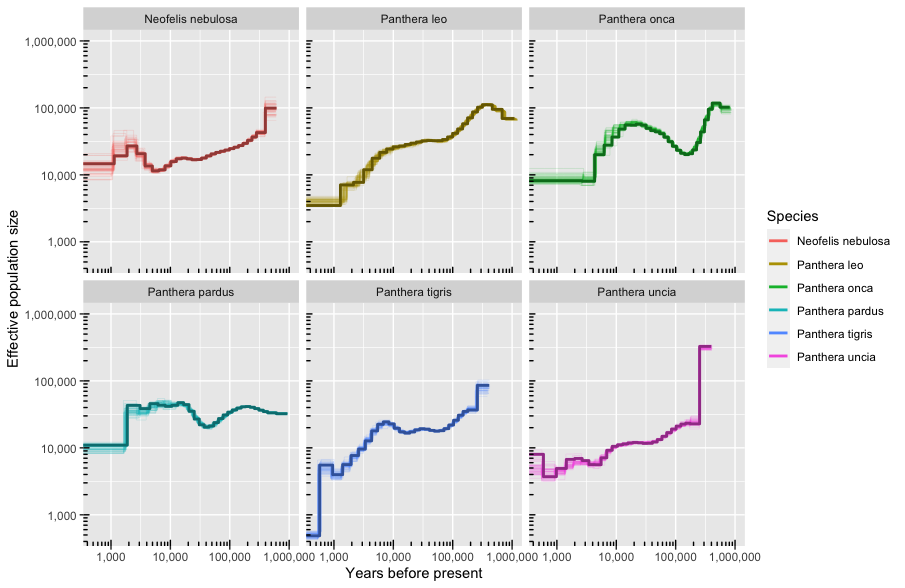

**Figure S7**: Bootstrap PSMC plots for new mutation rates.

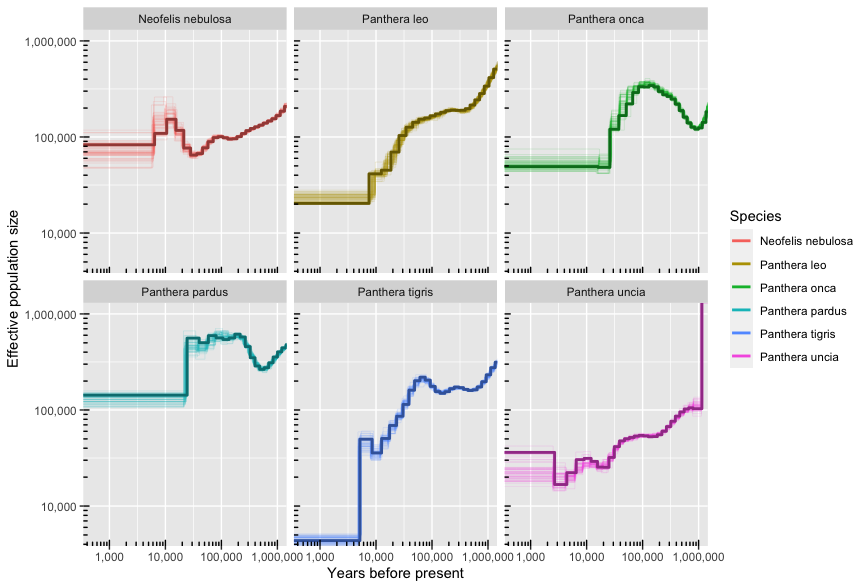

**Figure S8**: Bootstrap PSMC plots for previous minimum mutation rate.

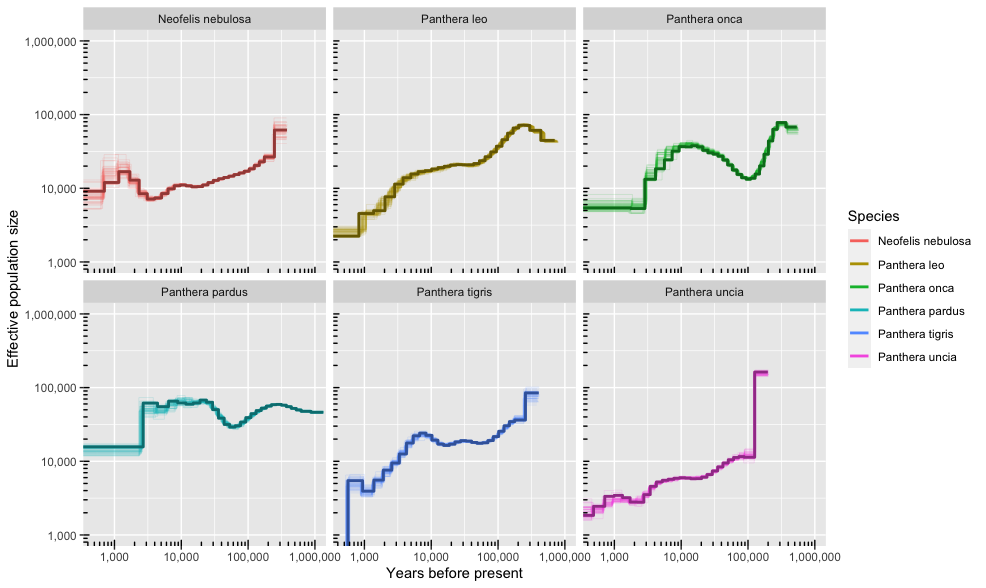

**Figure S9**: Bootstrap PSMC plots for previous maximum mutation rate.
